# Gene synthesis allows biologists to source genes from farther away in the tree of life

**DOI:** 10.1101/190868

**Authors:** Aditya M. Kunjapur, Philipp Pfingstag, Neil C. Thompson

**Affiliations:** Department of Chemical Engineering, Massachusetts Institute of Technology, Cambridge, MA, USA; Present Address: Department of Genetics, Harvard Medical School, Boston, MA, USA; Sloan School of Management, Massachusetts Institute of Technology, Cambridge, MA, USA; Present Address: School of Management, Technical University of Munich, Munich, Germany

## Abstract

Gene synthesis allows biologists and bioengineers to create novel genetic sequences and codon-optimize transgenes for heterologous expression. Because codon choice is key to gene expression and therefore cellular outcomes, it has been argued that gene synthesis will allow researchers to source genes from organisms that would otherwise have been incompatible, opening up more-distant parts of the tree of life as sources for transgenes. We test if this hypothesis is true for academic biological research using a 10-year data set from Addgene, the non-profit plasmid repository. We observe ~19,000 unique genes deposited to Addgene and classify them by whether they are natural or synthetic using a nucleotide-only technique that we develop here. We find that synthetic genes are an increasing share of Addgene deposits. Most importantly, we find direct evidence that researchers are using gene synthesis to source genes from more genetically-distant organisms, particularly for genes that are longer and thus might otherwise be particularly challenging to express. Thus, we provide the first empirical evidence that gene synthesis is leading biologists and bioengineers to sample more broadly across the rich genetic diversity of life, increasingly making that functionality available for industrial or biomedical advances.

## Introduction

Biologists and bioengineers work disproportionately with a small number of model organisms that are relevant for human health or industrial fermentation^1^. However, billions of years of evolution has provided a rich exploration of gene sequences across the full diversity of life. Researchers may wish to access and transfer these genes to test genetic hypotheses or to endow their favorite model organisms with novel traits or functionality^2,3^. In the first industrial example of recombinant DNA technology, Eli Lilly and Genentech expressed a synthetic gene encoding human insulin in the model bacterium *Escherichia coli* for drug manufacturing^4^. Soon afterwards, biologists began sourcing genes encoding thermostable polymerases^5^ from thermophilic bacteria and the well-known green fluorescent protein^6^ from the jellyfish as routine research tools. More recent biological research focused on mammalian models has featured considerable introduction of bacterial genes, notably the targeted genome editing tool CRISPR-Cas9^7–9^ and tools for optogenetics^10,11^. The growing field of synthetic biology also drives gene transfer because the genome sequences of non-model organisms present a treasure trove of potentially novel and orthogonal genes for testing in model organisms^12^.

Even though an increasing number of sequences are becoming accessible through next-generation genome and metagenome sequencing efforts^13^, many genes are expected to express poorly in heterologous organisms due to differences between the codon usage of the source organism and the new expression organism, differences in GC content, or the presence of expression-limiting regulatory elements within their coding sequence^14,15^. These concerns worsen as sequences get longer because the potential for problematic codons increases, as does the time required to manually convert these codons using PCR-based or restriction enzyme-based approaches. In contrast, gene synthesis can faithfully recode natural sequences of large lengths at once using synonymous codons that more closely reflect the expression organism and preserve the natural protein sequence^16^. Such codon-optimization is so valuable that commercial gene synthesis service providers typically offer this option by default.

Based on the improvement in heterologous gene expression expected from codon-optimization, we hypothesize that gene synthesis is accelerating the transfer of genes across the tree of life. Thus, we predict that the average genetic distance between source and expression organism will be greater for synthetic genes, compared to natural genes, and that this effect will be stronger for longer genes (which would otherwise be more difficult to adapt to the expression organism). Ideally, we would test this hypothesis directly on codon usage mismatch, but because codon usage tables are not available for many of the organisms that we study, we instead test our hypothesis using a correlate^17^: genetic distance.

The data for our analysis comes from the Addgene plasmid repository^18^. Because depositors to Addgene do not clearly indicate whether their deposited gene is natural or synthetic, we develop and test a method to determine this based only on the nucleotide sequence. To the best of our knowledge, no other bioinformatics approach fulfills this objective, so we anticipate that this method will be of independent interest to biotechnology companies and to intelligence agencies. Having classified the genes in the Addgene database as natural or synthetic, we examine how they are expressed and whether this is consistent with the hypothesized role of synthesis as allowing researchers to access genes from farther away in the tree of life.

The Addgene repository is a fruitful place for us to investigate these trends because it is one of the “go-to” repositories for academic access to plasmids^19^, and the most-used source for CRISPR-Cas9^20^. This prominence is evidenced in the rapid rise of the number of plasmids deposited over time (Fig. 1A), orders per year (Fig. 1B), and new labs depositing plasmids (Fig. 1C). Addgene is also used across a wide variety of expression platforms (Fig. 1D). These data demonstrate that Addgene is both a key plasmid repository and that it is used by biological researchers harnessing organisms from across the tree of life. From our analysis of sequences deposited on Addgene, we find that the average genetic distance between source and expression organism is greater for synthetic genes compared to natural genes and that this difference increases at longer sequence lengths.

**Figure 1.**
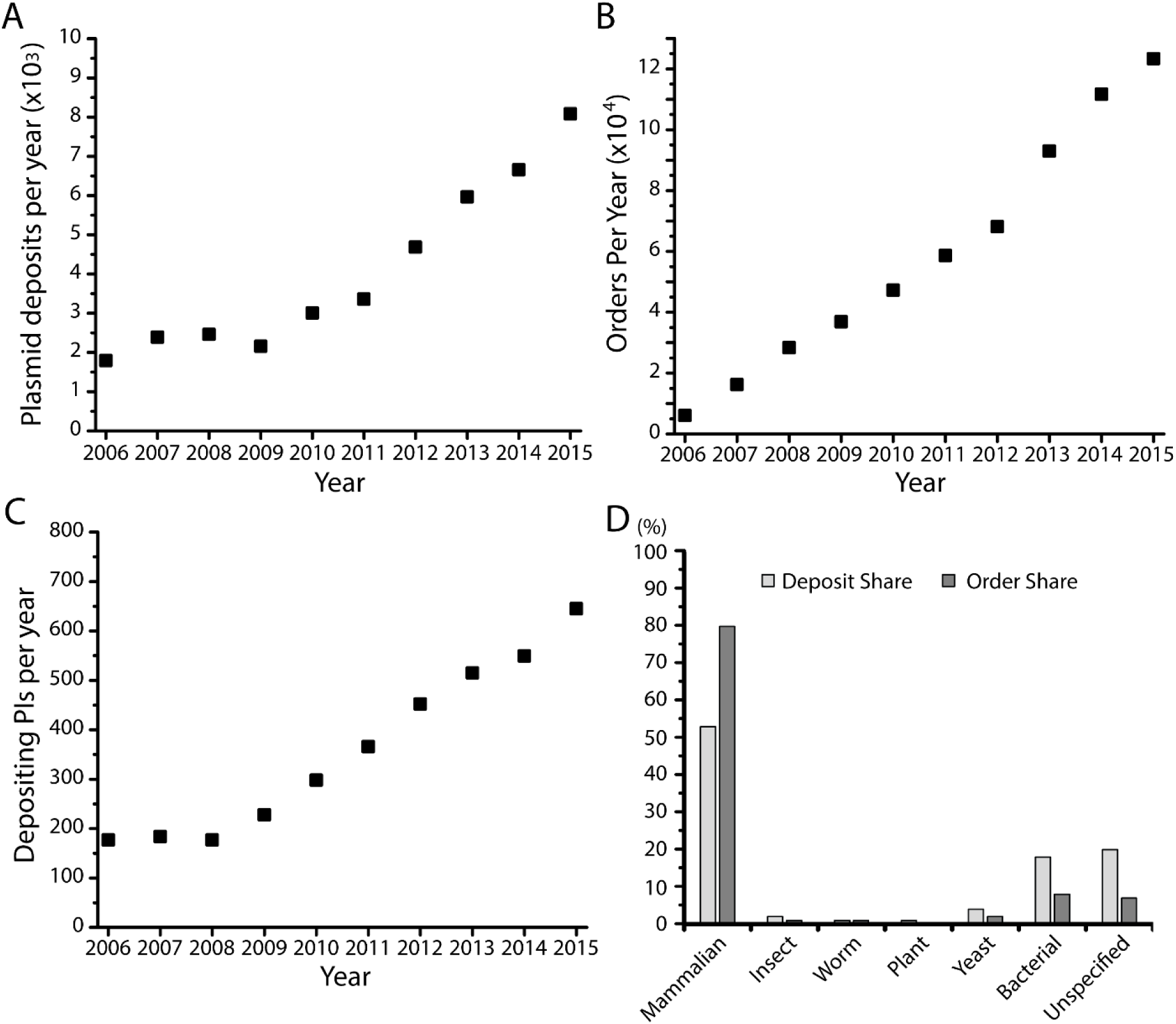
Descriptive statistics of the Addgene plasmid repository, a representative source of sequences used in academic biological research. (A) Number of plasmids deposited per year during 2005-2016 in Addgene. (B) Number of orders per year from Addgene. (C) Number of unique depositing principal investigators (PIs) per year. (D) Share of deposits and orders for plasmids across different expression platforms.

## Results

### Classification scheme for natural or synthetic genes

There are many plausible definitions that could be used for defining whether a sequence is “natural” or “synthetic.” We define a natural gene sequences as one that is found in naturally occurring genomes or metagenomes, including sequences that contain small deviations such as those resulting from natural evolution or from minor human interventions such as appending of short tags. We also consider cDNA sequences as natural given that they can be generated from naturally occurring mRNA using reverse transcription. In contrast, we apply the term synthetic to gene sequences that contain significant deviations from any single known contiguous naturally occurring sequence. Our definition is pragmatic and has limitations which reflect that we are only using the nucleotide sequence for the classification. For example, it is necessarily the case that if a researcher ordered a part from a gene synthesis company that was identical in every base to a natural sequence, we would classify that gene as natural.

To learn which attributes best predicted this classification, we considered two sets of attributes: “intrinsic” properties that we could determine from the sequence (such as GC content and rare codon percentage); or “comparative” properties that we could determine through similarity comparisons with a reference sequence database (such as query coverage – “QCov” – or percentage identity – “%Id”). To gather the comparative information, we used nucleotide BLAST^21,22^ to test each sequence against the NCBI RefSeq database, a comprehensive database of naturally occurring genomes, metagenomes, and cDNA libraries^23,24^ , and extracted comparison data for the best alignment entry (Fig. 2A). We consider the organism corresponding to the best alignment entry as the “source organism” for a gene sequence.

**Figure 2.**
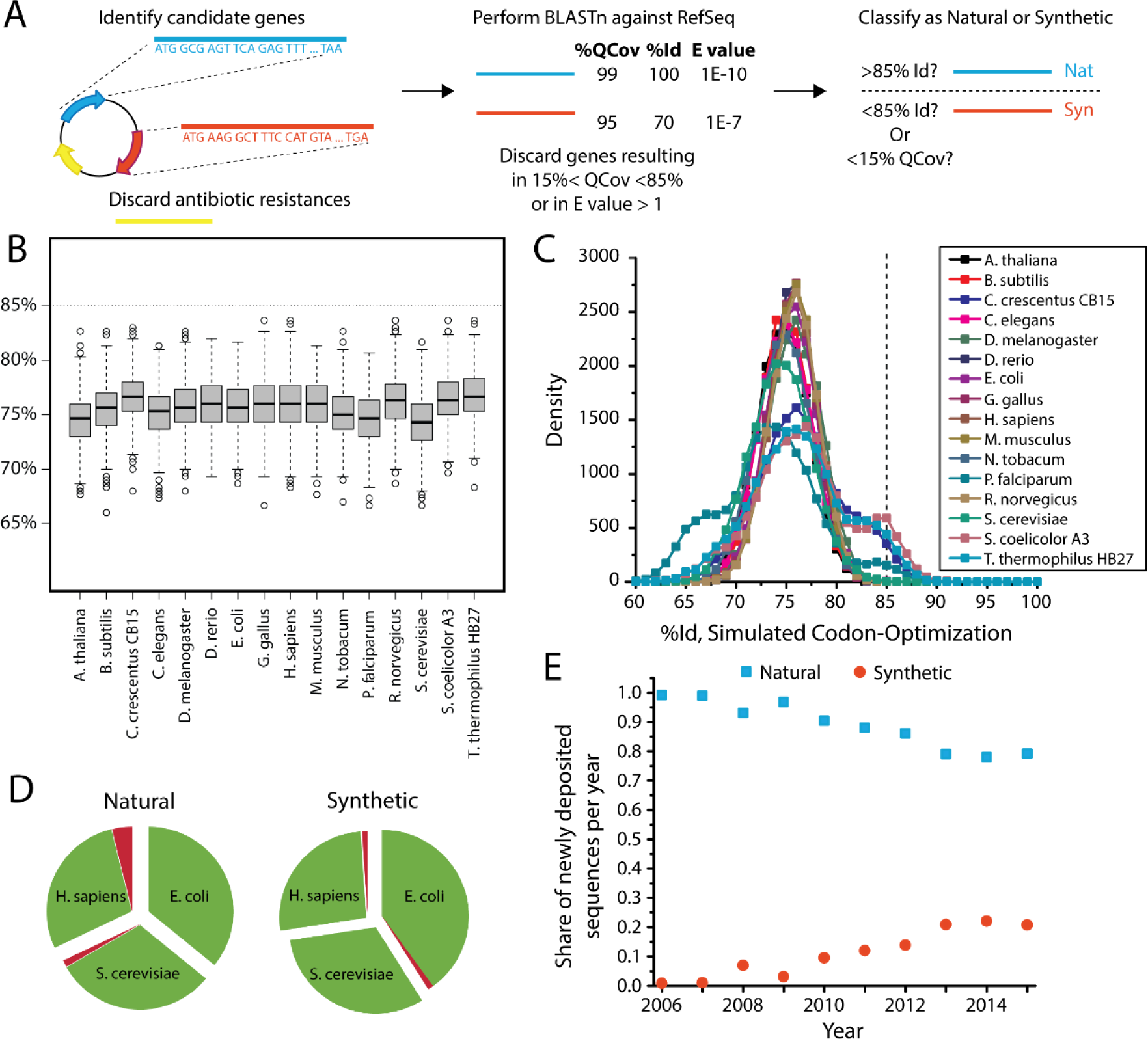
A nucleotide-based sequence classification scheme for determining natural and synthetic gene sequences. (A) Overall classification workflow. (B) Simulated sequence percentage identity (%Id) of a codon-optimized sequence originating from 16 organisms (including model organisms and organisms with distinct codon usage) for expression in *H. sapiens.* (C) Simulated %Id resulting from pair-wise codon-optimization of a hypothetical sequence for all 16 organisms and aggregating by expression organism (D) Results from application of classification scheme to a manually constructed test set of natural and synthetic gene sequences for expression in *Escherichia coli*, *Saccharomyces cerevisiae* (baker’s yeast), and *Homo sapiens*. Regions colored in green represent the proportion of sequences successfully identified by our classification scheme, and regions in red represent unsuccessful cases. (E) Share of natural and synthetic sequences deposited to the Addgene repository, as determined by application of classification scheme.

Because many published or publicly disclosed codon-optimization procedures use a weighted Monte Carlo approach proportional to codon abundance (or codon adaptation index, CAI)^25–29^, we surmised that there might be an effective cutoff value for %Id below which there would only be synthetic sequences. To quantify this theoretical cutoff, we pursued two strategies. First, we computed the average expected %Id of a nucleotide sequence assuming randomized codon-substitution *without* weighting by the codon usage of any particular organism. Each codon substitution thus produced an expected %Id based only on the number of codon possibilities for each amino acid and the nucleotide substitutions between them. Weighting these values by the amino acid occurrence frequency in nature^30^ indicates that a randomly codon-substituted sequence should, on average, have 75 %Id compared to the starting non-substituted sequence (Supplementary Tables 1-3). This provides the baseline against which we compare our second strategy, which tests the expected %Id from codon-optimization for specific organisms. We did stochastic simulations of all potential pairs of 16 different organisms, using their actual codon usage tables. On average, these simulations provided similar results, for example for expression of human sequences in other organisms (Fig. 2B) had an average of 75 %Id and all simulations fell below 85 %Id. Codon optimization across other organism pairs revealed important variation from the 75 %Id average: organisms with highly-dissimilar codon usage produced 65 %Id, whereas optimizing organisms with highly-concentrated codon usage back to themselves produced 85 %Id (overall distribution summarized in Fig. 2C, the ‘shoulders’ of which, at 65% and 85%, represent these extremes). Because model organisms contain more typical codon usage, transfers of genes between two organisms with extreme codon-usage are infrequent. Together our strategies provide strong evidence that Monte-Carlo based codon optimization methods are strongly likely to leave telltale signals in the %Id when compared back to the pre-codon-optimization sequence.

To determine which other attributes could inform our classification, we pursued a supervised learning approach. We constructed a training set consisting of 83 gene sequences populated with natural and synthetic genes for expression in *E. coli, Saccharomyces cerevisiae* (baker’s yeast), and *Homo sapiens* (Supplementary Tables 4-10) with details on: %QCov, %Id, GC content and percentage of rare codons used (2%, 5%, and 10%). We applied a “random forest” approach to this training set and determined that sequence %Id below 85% was the best predictor of it being synthetic, aligning well with our theoretical results. Using this classification criteria on a test set of 173 manually identified sequences, we observed over 97% success (Fig. 2D). This aligns well with our theoretical simulation results, which predict that 98.6% of synthesized sequences will lie below the 85 %Id threshold. To limit our analysis to non-fusion proteins, and to be consistent with the %Id threshold, we also impose an 85 %QCov threshold, which our random forest approach deems an equally strong predictor.

### Application of classification scheme to the Addgene database

We applied our BLAST-based classification scheme to the 19,334 unique genes contained in the Addgene database from 2006-2015 to determine which were synthetic and which were natural. For this analysis, we excluded known antibiotic resistances and those sequences that appeared to encode fusion proteins, based on their BLAST results (see Supplementary Methods). We find that the share of synthetic gene sequences deposited in Addgene has increased over time (Fig. 2F). By 2015, synthetic sequences made up over 20% of the genes in newly deposited plasmids, up from less than 1% in 2006. The increasing abundance of synthetic sequences is consistent with the order of magnitude decrease in the cost of gene synthesis over this period^31–33^.

### Examination of differential transgene expression by taxonomic grouping

Using our classification and BLAST results, we investigated patterns of source and expression organisms for natural and synthetic gene sequences. Because Addgene expression fields contained terms broader than specific organisms, we grouped expression into six categories: Mammalian, Worm, Insect, Plant, Yeast, and Bacteria (Supplementary Table 11). For source organisms, we used NCBI taxonomic classification of the best alignment organism to categorize at multiple levels spanning phylum to individual species. Many synthetic sequences resulted in no BLAST alignment to any sequence in the RefSeq database, and these were designated as “No Hit.” Sequences that result in “No Hit” are likely to be *de novo* synthetic sequences that deviate from any known protein. We report the proportion of sequences that fit into this category and where they are expressed because this is of independent interest, but we ignore these sequences for subsequent genetic distance calculations since they lack a source organism. Additionally, for sequences that best aligned to viral sequences, we included a “Virus” category that exists outside of taxonomic relationships for living organisms. We binned all sequences with available source organisms by phylum in accordance with common taxonomic practice, reflecting the relative genetic homogeneity within phyla

Estimating genetic distance between source and expression categories is not possible using existing taxonomic systems because they are not quantitative and because such comparisons cannot be made at the phylum level. Instead we measure distance using 16S or 18S ribosomal RNA (rRNA) sequence from the SILVA database^34^. rRNA is highly evolutionarily conserved and can function as an evolutionary chronometer since 18S rRNA is the eukaryotic nuclear homolog of 16S rRNA in prokaryotes^35,36^. We then constructed a phylogenetic tree using the web tool Phylogeny.fr^37^, and extracted genetic distances for each source-expression pair based on the most-common organism in that phylum in the Addgene database (Fig. 3A and Supplementary Fig. 1). These genetic distances represent the fraction of mismatches at aligned positions, as is conventional in phylogenetic analysis.

**Figure 3.**
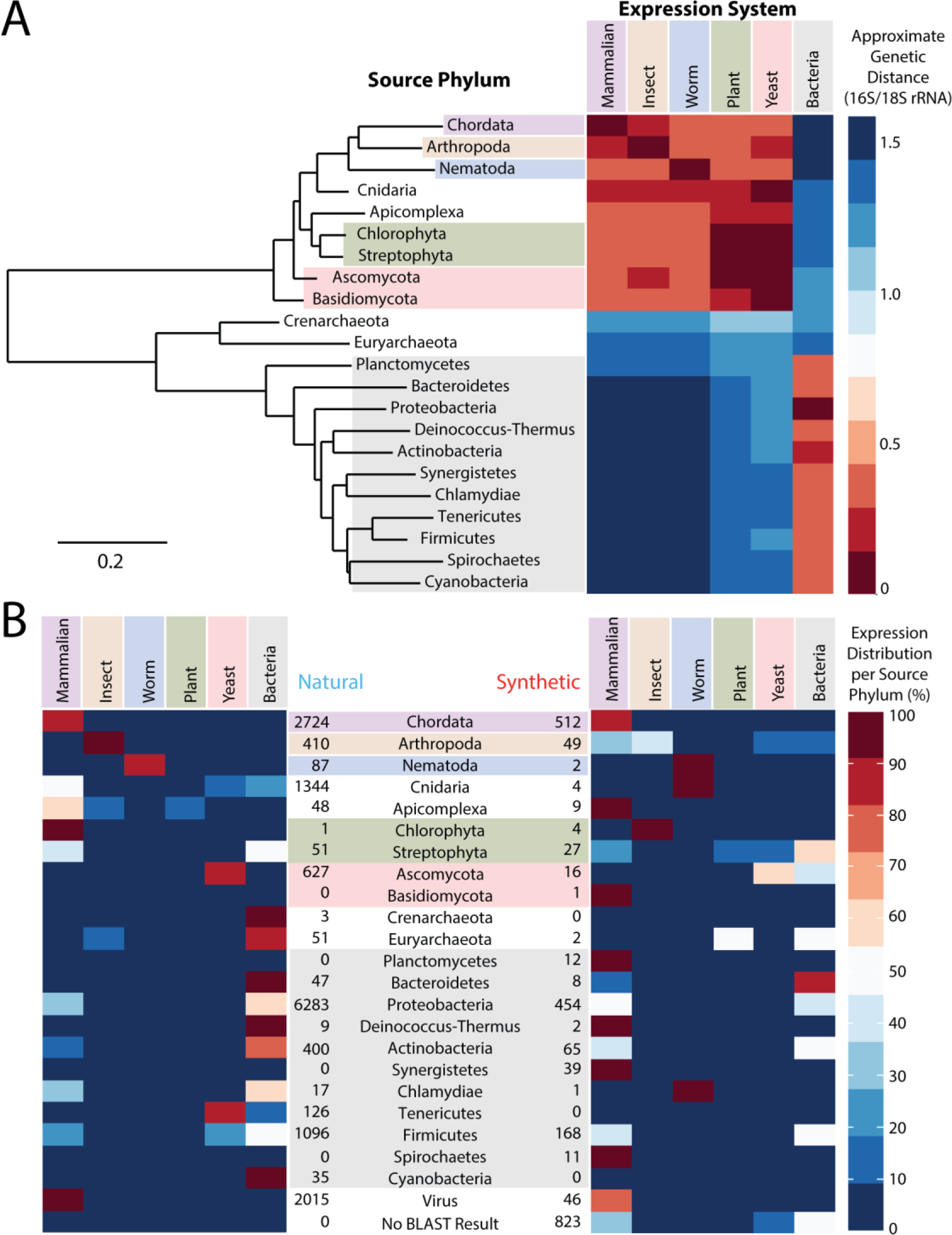
Mapping of genetic distances and transgene expression for Addgene sequences. (A) Phylogenetic tree of the most common source phyla and corresponding heatmap displaying genetic distance of different expression platforms. (B) Heatmaps displaying source phylum and expression platform for (left) natural and (right) synthetic genes.

We display heatmaps showing the number of natural and synthetic gene sequences in the Addgene database corresponding to a source-expression category pairs across the 22 most common phyla (Fig. 3B). From these heatmaps, we can make several observations spanning the relative magnitude of phylum sourcing, the kinds of gene transfers occurring, and the differences in these aspects between natural and synthetic genes. Though the most common expression system for Addgene plasmids is Mammalian, the largest source of unique gene sequences (by a significant margin) based on nucleotide BLAST is Phylum Proteobacteria. The next largest sources of unique gene sequences are Phylum Chordata, viruses, and Phylum Cnidaria, respectively. This may reflect the relative focus on studying vertebrate and viral genes as well as the importance of GFP in biological research. Sequences sourced from Proteobacteria are used at approximately similar levels in Mammalian and Bacterial expression systems, regardless of whether the sequence is identified as natural or synthetic. On the other hand, sequences sourced from Chordata are predominantly used only in Mammalian expression systems, regardless of whether the sequence is identified as natural or synthetic.

These heatmaps demonstrate significant transgene expression for both natural and synthetic gene sequences. The most frequent type of transfer is from Phylum Proteobacteria to Mammalian expression. Though this may be consistent with the predominance of deposits and orders of mammalian expression constructs from Addgene, it is striking that the frequency of transfer from Phylum Chordata to bacterial expression (essentially the reverse phenomenon) is far lower. A higher-level pattern observable in the heatmaps of Fig. 3B is their relationship with genetic distance shown in Fig. 3A. If genes were being most commonly expressed in their source organism, one would observe hotspots in Fig. 3B along a diagonal axis roughly from upper-left to lower-right. These hotspots are clear for animal expression platforms for both natural and synthetic genes. For natural genes, the pattern extends into many bacterial sequences (hotspots on the lower-right). However, for synthetic genes there is a marked change in the trend for bacterially sourced sequences. Hotspots frequently appear on the lower-left, indicating a high-frequency of mammalian expression of bacterially derived, synthetic sequences.

### Comparison of genetic distances between source and expression organisms for natural and synthetic genes

From these heatmaps it is difficult to quantify the differences in expression of natural and synthetic genes. Thus, we calculated genetic distances between the source and expression organism for each sequence. Overall, consistent with our main hypothesis, we find that the average genetic distance between source and expression organisms is greater for synthetic than for natural gene sequences, and that this distinction is highly statistically significant. Table 1 shows these results through a series of regression specifications. In all cases, *Synthetic* is a binary variable which is 1 if our classification system deems that sequence synthetic, and 0 otherwise. Specification (1) shows that expression with synthetic sequences is, on average, 0.077 units (p-value <0.01) farther from the source organism than are natural sequences. Specification (2) shows that the gap between the genetic distance between synthetic and natural sequence use grows with sequence length, with each extra kilobase adding 0.117 units (p-value <0.01) to the difference. Specifications (3) and (4) confirm the finding of specification (2), but use the alternative dependent variable *Cross Kingdom*, which is a binary variable equal to 1 if the expression is cross-kingdom and 0 otherwise. These trends remain even if CRISPR-Cas9 sequences are excluded from the analysis (Supplementary Table 12). Fig. 4 uses a non-parametric local regression (loess) to show the relationship between gene length and genetic distance, for both natural and synthetic sequences. The shade regions represent one standard error.

**Table 1:** Regression results comparing natural and synthetic sequences from the Addgene Database

|  | Dependent variable: |  |  |  |
| --- | --- | --- | --- | --- |
|  | Genetic Distance | Genetic Distance | Cross Kingdom | Cross Kingdom |
|  | OLS<br>(1) | OLS<br>(2) | OLS<br>(3) | Logit<br>(4) |
| **Constant** | 0.499***<br>(0.006) | 0.606***<br>(0.009) | 0.572***<br>(0.007) | 0.452***<br>(0.035) |
| **Synthetic** | 0.077***<br>(0.018) | -0.041<br>(0.029) | -0.017<br>(0.022) | -0.228**<br>(0.093) |
| **Gene Length [kb]** |  | -0.112***<br>(0.007) | -0.098***<br>(0.005) | -0.587***<br>(0.035) |
| **Gene Length [kb] \* Synthetic** |  | **0.117\*\*\***<br>**(0.013)** | **0.076\*\*\***<br>**(0.010)** | **0.495\*\*\***<br>**(0.049)** |
| Observations | 14,745 | 14,745 | 14,745 | 14,745 |
| R2 | 0.001 | 0.018 | 0.023 |  |
| Adjusted R2 | 0.001 | 0.018 | 0.023 |  |
| Log Likelihood |  |  |  | -10,000.32 |
| F statistic | 17.82 | 91.8 | 115.72 |  |

\*p<0.1; \*\*p<0.05; \*\*\*p<0.01

**Figure 4.**
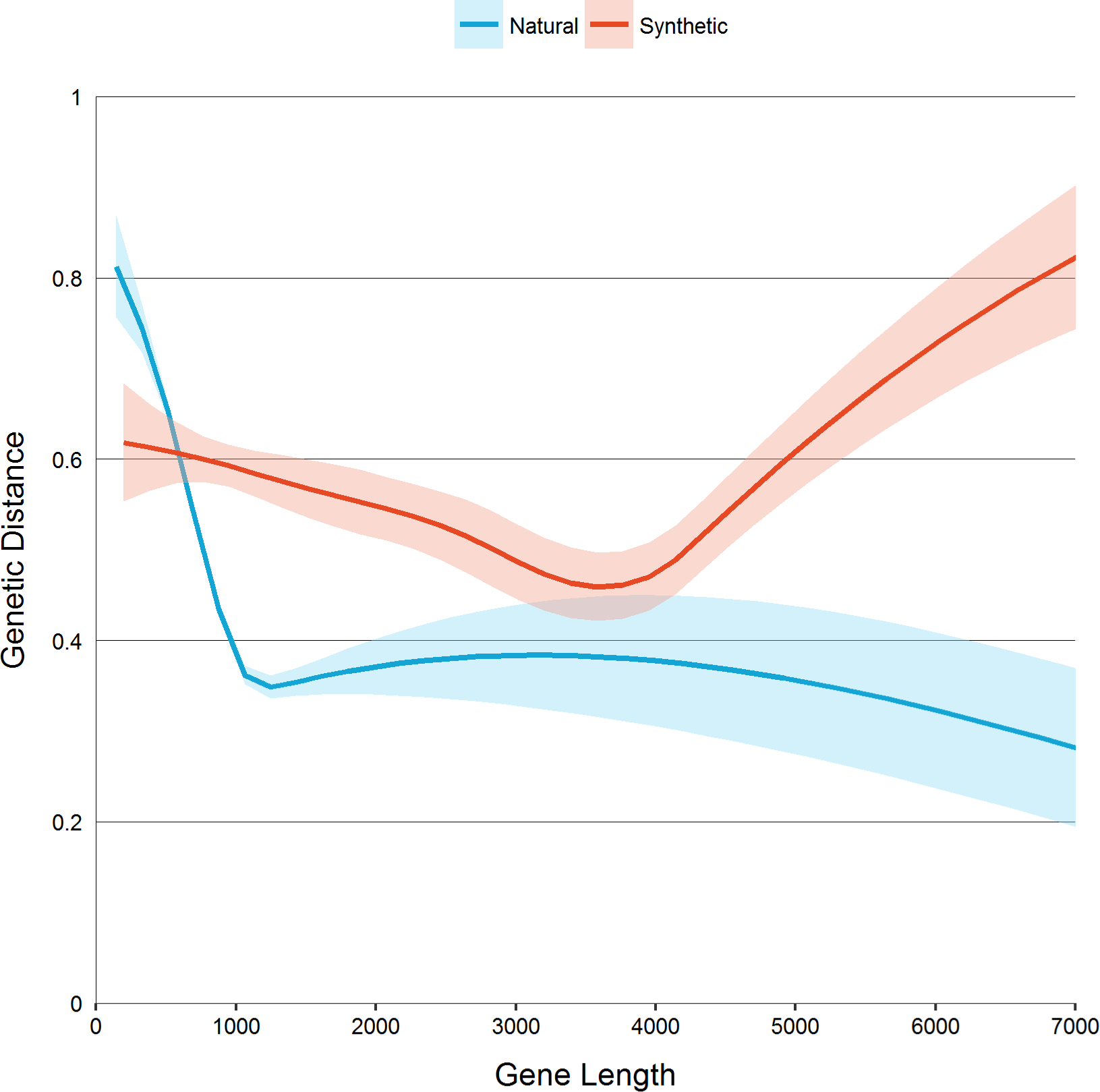
Local regressions (LOESS) of genetic distance and gene length for natural and synthetic sequences in the Addgene database.

Our results suggest that the longer a *natural* gene sequence is, the less likely it is to be transferred into another organism by researchers. This observation is consistent with the perception that longer unmodified gene sequences are generally more difficult to express and that, as sequence length grows, so does the likelihood that there will be sequence regions that are troublesome to express in another organism. In contrast, synthetic sequences experience little to no drop in genetic distance as gene length grows, and at large lengths are used predominantly for transfer across distant organisms. Thus we conclude that gene synthesis enables transgene expression at a much higher rate than traditional techniques, and that this difference is both scientifically and statistically significant.

## Discussion

This paper develops, to our knowledge, the first nucleotide-only-based method for determining whether a genetic sequence is synthetic or natural. Grounded both in theory and empirical testing, we find that we can correctly predict with 97% accuracy on a novel data set. This is important not only for our project, but as a starting place for those in the larger genetics community that are interested in understanding the provenance of a genetic sequence or in surveillance.

Using our classification method, we provide the first empirical evidence that gene synthesis is being widely used by practitioners to source genes from genetically-distant organisms. These synthesized parts are not only notably more distant from their source organism than are natural sequences, but this gap grows as sequence length increases. The importance of gene synthesis in accessing this treasure trove of genetic diversity should not be undervalued. Previously researchers could look for parts in a narrow genetic neighborhood where transfer would be relatively easy, or source more broadly at the cost of potentially incurring much greater engineering effort with little guarantees of eventual success. Today those same researchers can source their parts wherever makes the most biological sense with the knowledge that gene synthesis will overcome one of the main expression challenges for them. As such, gene synthesis is allowing biologists to source genes from farther away in the tree of life. This promises to be of scientific, industrial and medical use, to the great benefit of biologists and society at large.

## Acknowledgements

We are tremendously indebted to Addgene for sharing their data, answering questions as they arose, and providing manuscript feedback. We thank Dr. Alec Nielsen (MIT), Dr. Darrell Ricke (MIT), and Dr. James Comolli (MIT) for discussions about BLAST. We are grateful to Dr. Nili Ostrov (Harvard) and George Chao (Harvard) for discussions about genetic distance calculations. AMK was supported by a National Science Foundation Graduate Research Fellowship PP was supported by two grants of the European Union's Seventh Framework Programme FP7. The collaborative research project ST-FLOW (KBBE-2011-5 – Grant Agreement number 289326) and the People Programme (Marie Skłodowska-Curie Actions – Grant Agreement number 612614) and NCT was supported by a grant from MIT.

**Online Methods** and **Supplementary Information** included separately.

